# Much ado about nothing: Modelling amino acid replacement with predicted protein structures

**DOI:** 10.64898/2025.12.05.692347

**Authors:** Lukas Buschmann, Sarah Naomi Bolz, Ferras el-Hendi, Negin Malekian, Michael Schroeder

## Abstract

Substitution matrices like BLOSUM62 model the likelihood of replacement of amino acids in evolution. Substitution matrices are used in protein sequence alignment tasks. Since the introduction of BLOSUM62 over three decades ago, many matrices have been released.

Yet, to date, no effort uses large amounts of 3D structures predicted by AlphaFold. Here, we define AFSM, the AlphaFold Substitution Matrix derived from over 20,000 predicted 3D structures following the BLOSUM methodology. We benchmark AFSM against BLOSUM62 and 16 other matrices on five tasks in multiple sequence alignment (MSA) and protein homology search. Our analysis surprisingly reveals that all matrices perform similarly. Only when there are few sequences in an MSA, then BLOSUM62 and AFSM perform better than using no matrix. This suggests that substitution matrices were most beneficial when there was little sequence data. We corroborate this argument by showing that embeddings, which are computed from billions of sequences, perform better than substitution matrices, when sequence data is sparse. Taken together, this suggests that structural data does not improve BLOSUM62. But increased sequence data makes extrapolation with substitution matrices obsolete. Nonetheless, BLOSUM62 continues to capture chemists’ intuition on amino acids by providing numerical values implicitly reflecting physicochemical properties, and it remains indispensable for direct comparison of two sequences.

## Introduction

Substitution matrices quantify the likelihood of amino acid replacements in evolution. They have been fundamental in the construction of multiple sequence alignments (MSAs) and remote homology search. They guide the alignment process by assigning scores that favor biologically meaningful matches and penalize unlikely substitutions, thereby enabling the detection of evolutionary relationships between sequences. Two widely used matrices, PAM and BLOSUM (Dayhoff et al. 1978), date back to 1978 and 1992, a period in which the amount of sequence data rose from hundreds to a hundred thousand. This unprecedented growth continued, with a hundred million sequences available in 2009 (www.ncbi.nlm.nih.gov/genbank/statistics). As sequence data grew, new substitution matrices were published. PAM was refined with more sequence data by Gonnet et al. (Gonnet et al. 1994) and an improved statistical model (Müller et al. 2000). For BLOSUM, the re-implementations RBLOSUM and CORBLOSUM (Hamacher et al. 2016) were published.

In parallel, in 2000, the first structural approaches emerged. Protein structure is more conserved than sequence (Illergård et al. 2009), which implies that it should be better suited for the construction of matrices. However, there was far less structural data available than sequence data. In 2000, when there were some ten million sequences, the protein data bank PDB held 13,583 protein structures (RCSB.org, Bernstein et al. 1977). Far less, but sufficient for Prlić et al. to build two substitution matrices, SDM and HSDM (Prlić et al. 2000). Two decades later, PDB held nearly ten times as much data, and two more approaches used structural data. Jia et al. (Jia et al. 2021) analyzed co-evolving residues in over 5,000 structural families (PROTSUB), and Keul et al. (Keul et al. 2017) built their PFASUM matrix from Pfam (Bateman et al. 2021) seed alignments, which in turn use structural data.

All of these matrices predate the arrival of AlphaFold (Jumper et al. 2021) and the availability of hundreds of millions of precomputed predicted protein structures (Varadi et al. 2022). Thus, in this paper, we introduce AFSM (see Figure 1), the AlphaFold substitution matrix. We follow the BLOSUM approach in construction, but derive substitutions from over 660,000 structural alignments for over 20,000 proteins. We then focus on how AFSM relates to BLOSUM62 and all of the above substitution matrices. Next, we introduce five benchmarks to evaluate the performance of AFSM. Two of the benchmarks address the construction of MSAs. The first, BAliBase (Thompson et al. 2011), is a standard in the field. It is well curated, but comparatively small. Therefore, we consider a second MSA benchmark derived from Pfam, which covers 21,000 protein families. We systematically compare the two benchmarks in terms of insertions, deletions, mismatches, matches, length of sequences, and number of sequences to assess their degree of difficulty. The MSA benchmark is complemented by benchmarks in remote homology search with varying degrees of sequence redundancy. The main result of this article is the systematic comparison of AFSM and all the above substitution matrices on these five benchmarks, revealing how they perform in identical setups.

**Figure 1:**
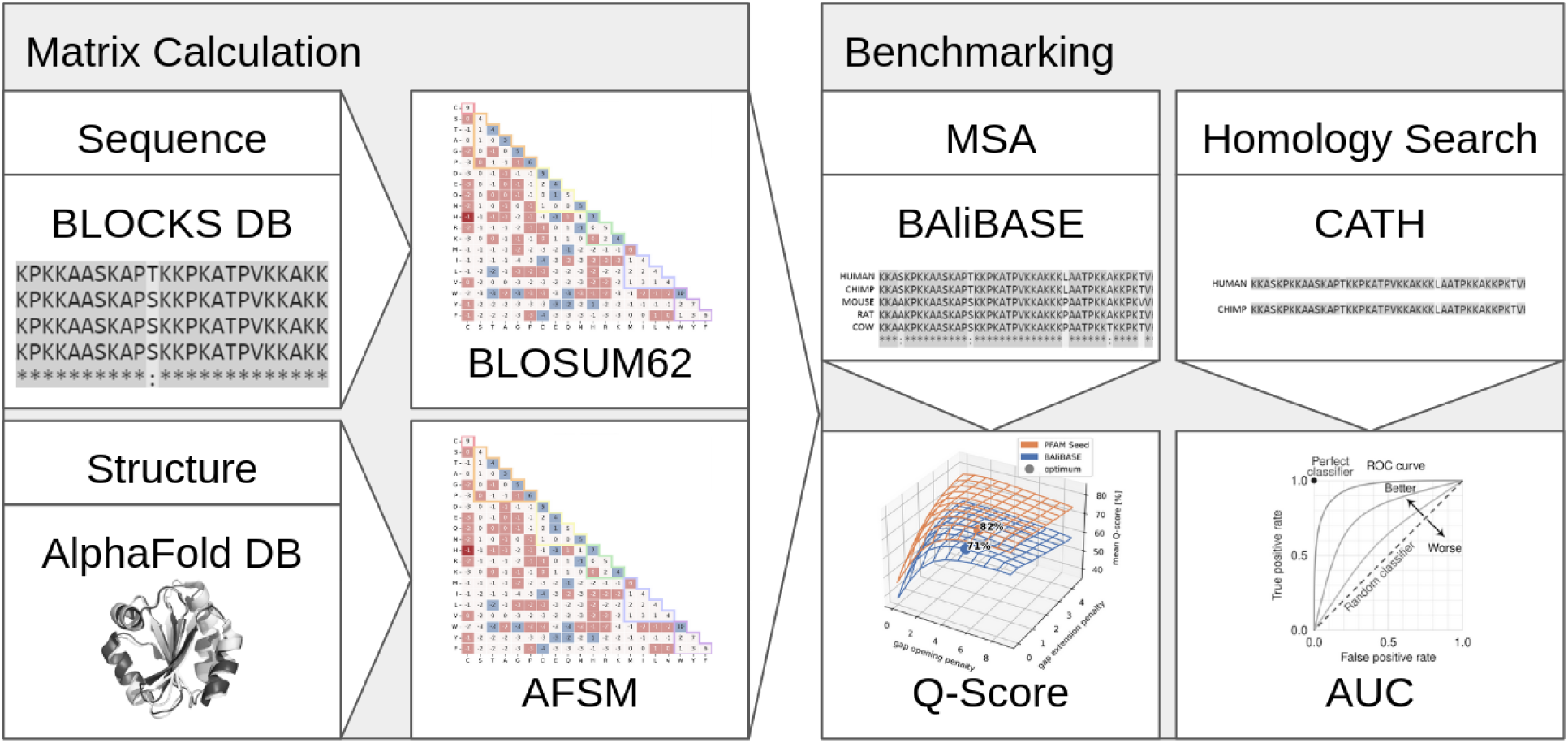
Graphical abstract

## Results

BAliBASE R10

Pfam Seed

### Construction of the AlphaFold Substitution Matrix (AFSM)

To construct the AlphaFold Substitution Matrix, we randomly selected 300 out of over 85,000 InterPro (Bateman et al. 2025) protein families. From each family, up to 100 protein sequences were randomly sampled, resulting in a dataset comprising approximately 21,000 unique proteins. For each of these proteins, we retrieved the predicted structures from the AlphaFold database (Varadi et al. 2022). For 3 of the 300 families, there were no structures at all, so that the subsequent analysis was based on 297 families only. Within each family, all possible protein pairs were generated. To mitigate redundancy and avoid over representation of highly similar sequences, we filtered out pairs with high sequence identity (>62%). The remaining pairs were subjected to structural alignment. Residue-residue correspondences were then extracted from structurally aligned regions, based on the spatial proximity of residues, resulting in ungapped pairwise alignments. To account for family size variability, alignments were weighted inversely proportional to the number of pairs per family, thus reducing bias from larger families. Finally, substitution scores were computed as log-odds in half-bit units, following BLOSUM. These scores were subsequently rounded to the nearest integer to produce the final matrix, as depicted in Figure 2A.

**Figure 2:**
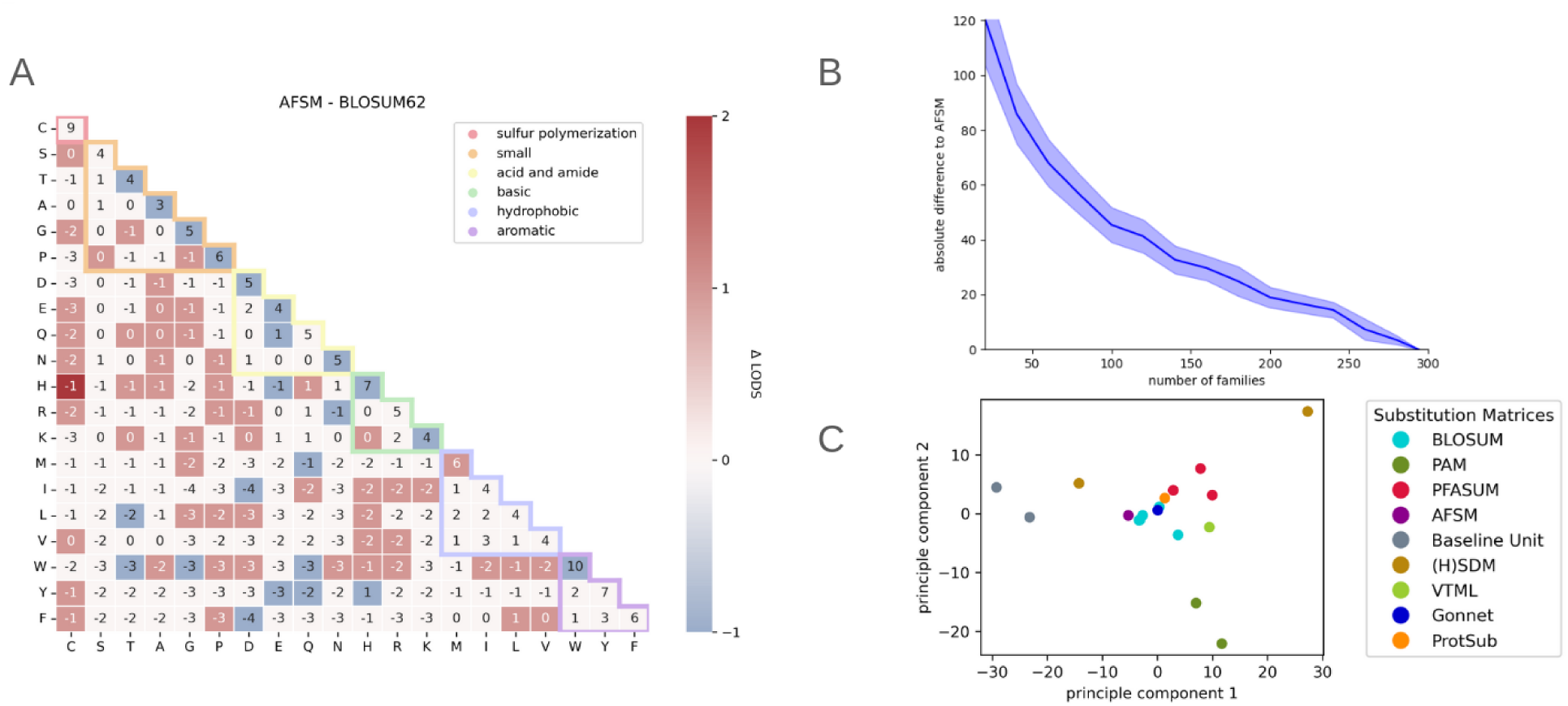
A. AFSM substitution matrix. Deviation from BLOSUM62 indicated by background color. Nearly half of the entries differ slightly. B Convergence behaviour in matrix construction. With an increasing number of families covered, AFSM construction quickly converges to the final matrix. Shown are the mean and one standard deviation across 1,000 random subsets for each subset size. C. Principal component analysis of 18 substitution matrices; from BLOSUM series: 45, 62, 80, CORBLOSUM61, RBLOSUM64; PAM series: 120, 250; PFASUM series: 31, 43, 60; (H)SDM: SDM, HSDM; Baseline Unit: 5/0, 5/-1.

To understand how the matrix depends on the amount of data, we varied the number of families. Figure 2B suggests that further increasing the number of families would have had a limited effect on the final AFSM matrix (using 200 instead of 297 families leads to a total difference of 20 only).

### Comparison of AFSM to BLOSUM62

Consider Table 1, which lists for BLOSUM62, AFSM, and selected other matrices their benchmark performance as well as meta information such as year of introduction, amount and type of data used, and similarity to BLOSUM62. Before presenting the benchmark results for all matrices in the table, let us focus on AFSM and BLOSUM62. Although AFSM uses structure instead of sequence data, and although the amount of data is over 100 times larger than for BLOSUM62, the AFSM matrix is similar to BLOSUM62, with 48% of 200 substitution scores identical. The 52% differing entries mostly show only small differences of ±1, except histidine and cysteine with a difference of 2 (see Figure 2A). Overall, AFSM has a very strong Pearson correlation of 96% to BLOSUM62. This strong resemblance is also reflected in Figure 2C, which depicts each substitution matrix listed in Table 1 as a high-dimensional vector, whose dimensions are reduced to two principal components (PCA). An interesting detail regarding AFSM and BLOSUM62 concerns their evolutionary distance (column Evo Dist in Table 1), which we captured as an absolute value of a ratio between average diagonal scores (match) over average off-diagonal scores (mismatch). BLOSUM45, 62, and 80 have distances of 5.3, 4.1, and 3.1, respectively, reflecting the degree of conservation of the underlying sequence data. With an evolutionary distance of 4.4, AFSM is closest to BLOSUM62 in this respect.

**Table 1.**
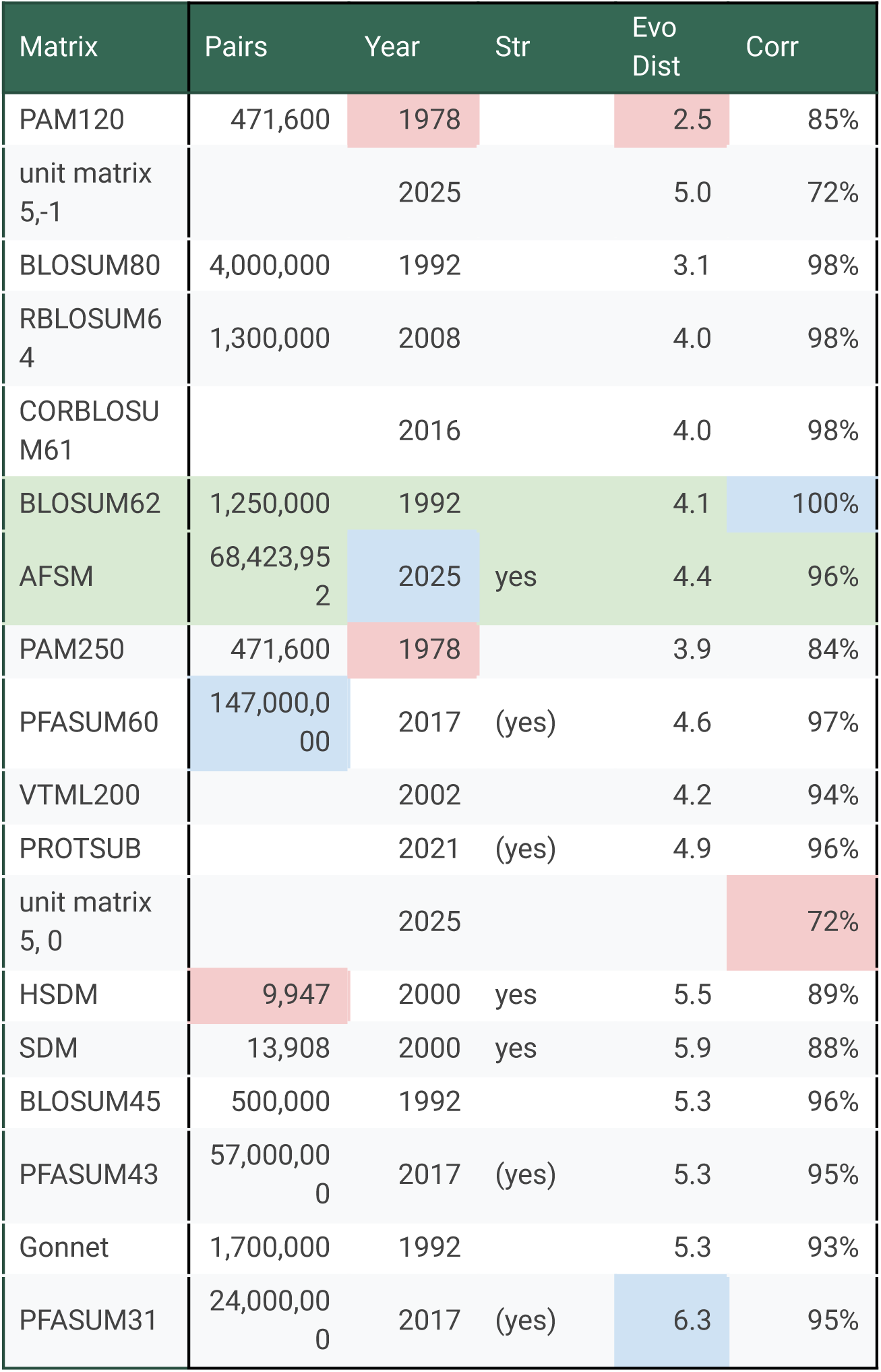
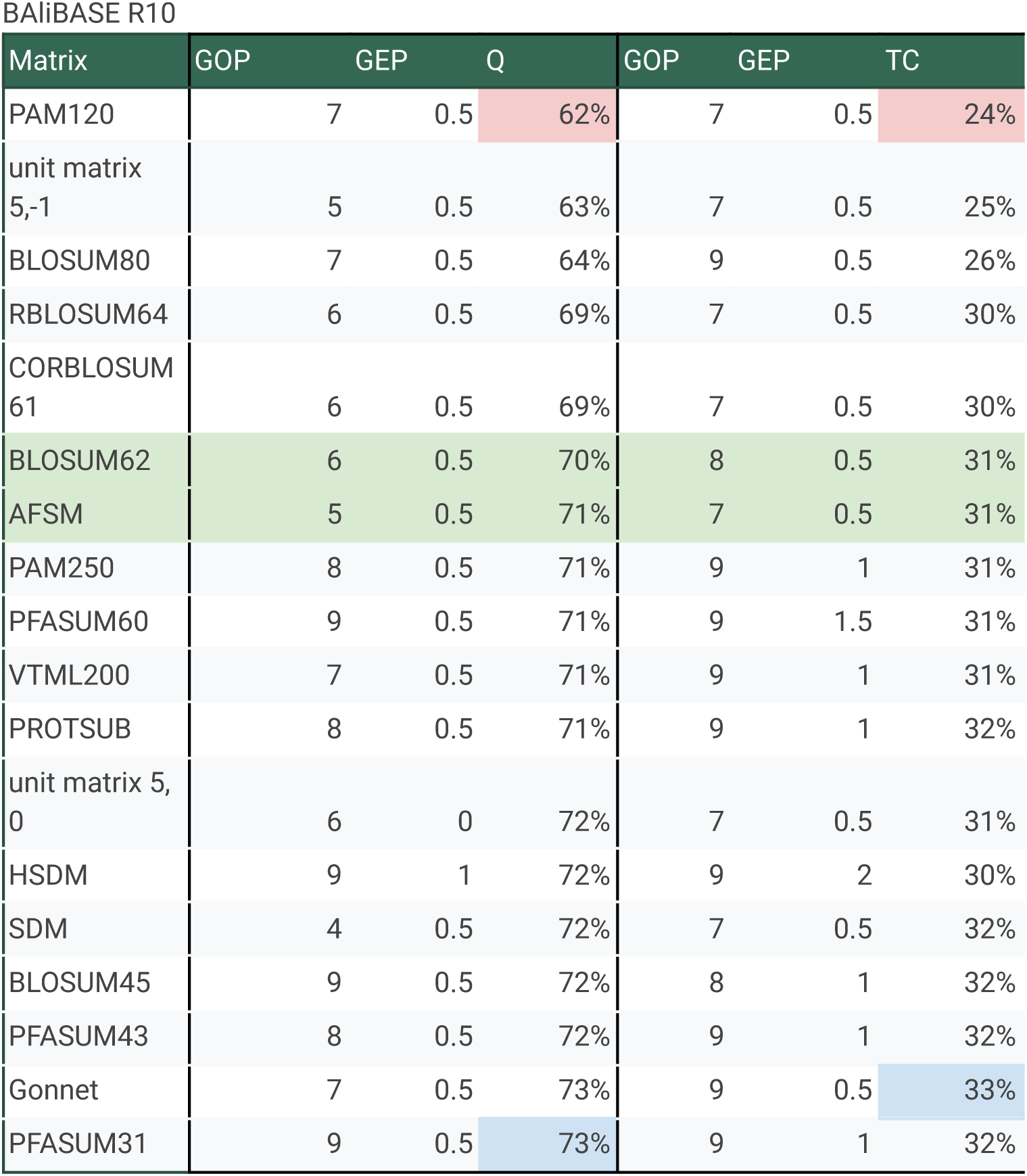
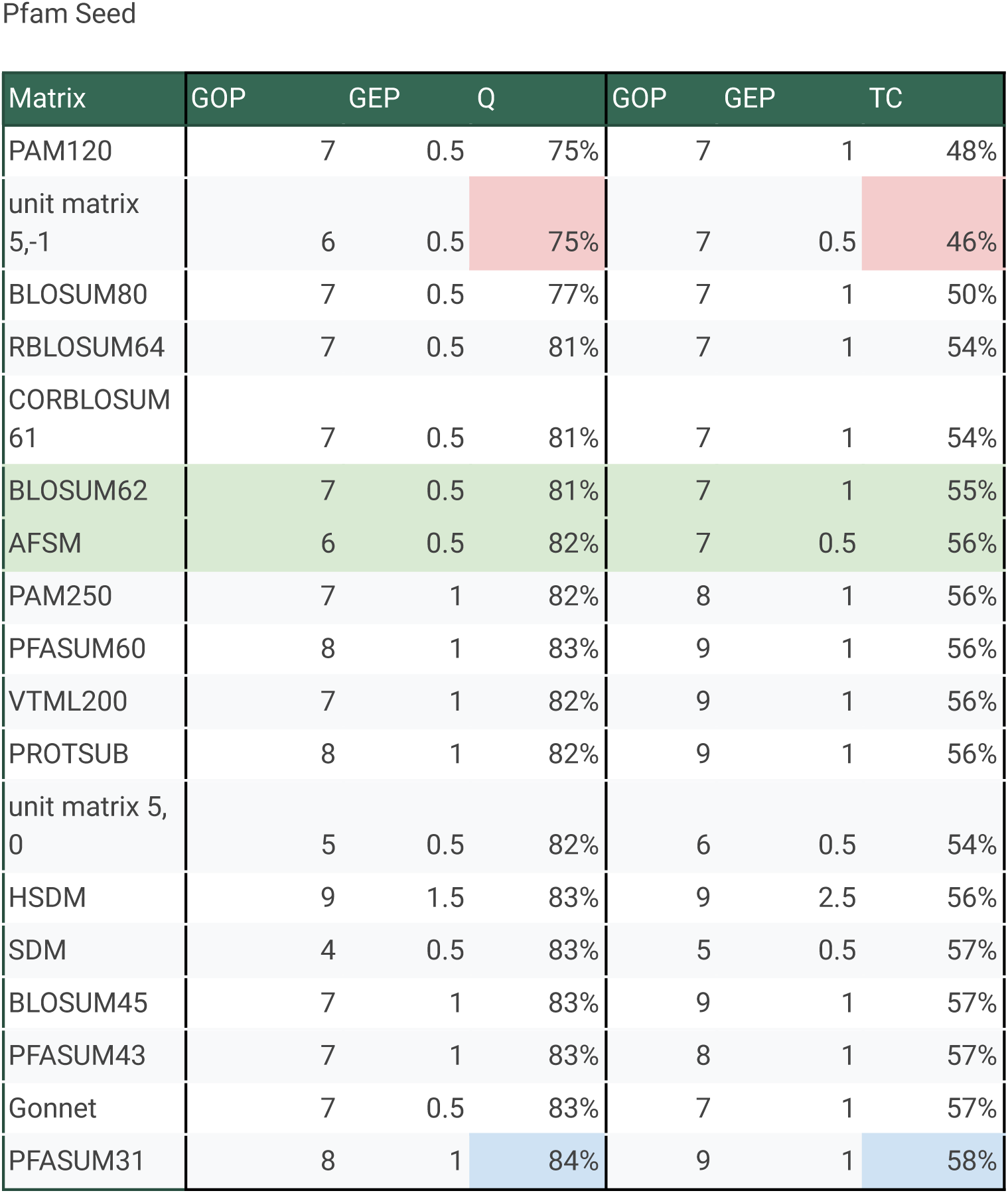
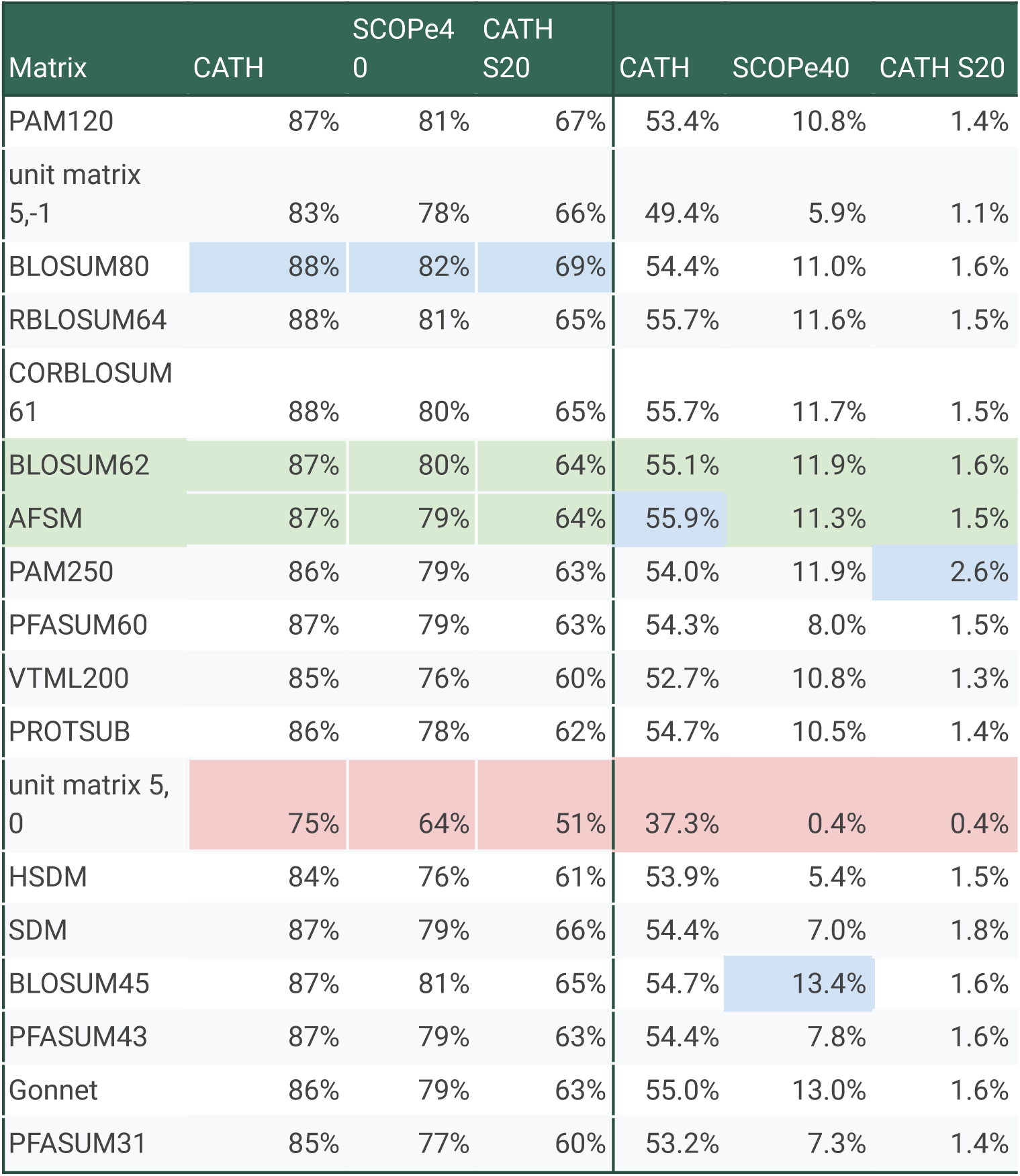
Overview of substitution matrices (in alphabetic order), their characteristics, and performance across benchmark tasks. AFSM and BLOSUM62 are highlighted as green rows. Per column, highest and lowest values are highlighted in blue and red, respectively. Pairs = estimated number of residue pairs used for construction. Year = year of publication. Str = structural data used. Evo Dist = evolutionary distance as absolute value of mean diagonal to off-diagonal; high = large, low = small evolutionary distance. Corr = Pearson correlation with BLOSUM62. GOP = gap opening penalty. GEP = gap extension penalty. Q=Q-score. TC=TC-score. CATH, SCOPe40, and CATH-S20 = AUC scores for corresponding task.

### Benchmarks

To evaluate the performance of AFSM, BLOSUM62, and the other substitution matrices, we employ two tasks: MSA construction and protein homology search. A widely used (Edgar 2009, Long et al. 2016, Keul et al. 2017) standard benchmark in MSA construction is BAliBase, which provides curated reference MSAs designed to represent a broad range of biologically relevant alignment scenarios. Since BAliBase R10 is comparatively small (209 MSAs), we added Pfam Seed alignments (18,061 MSAs) as a second benchmark. The Pfam Seed dataset offers a large collection of manually curated alignments across a diverse range of protein families. This inclusion allowed us to test the generalizability of AFSM on a broader and more diverse alignment set.

Despite their different scopes, the two benchmarks exhibit broadly comparable statistical properties relevant to MSAs (see Figure 3). However, there are notable differences in key factors that affect alignment difficulty.

**Figure 3:**
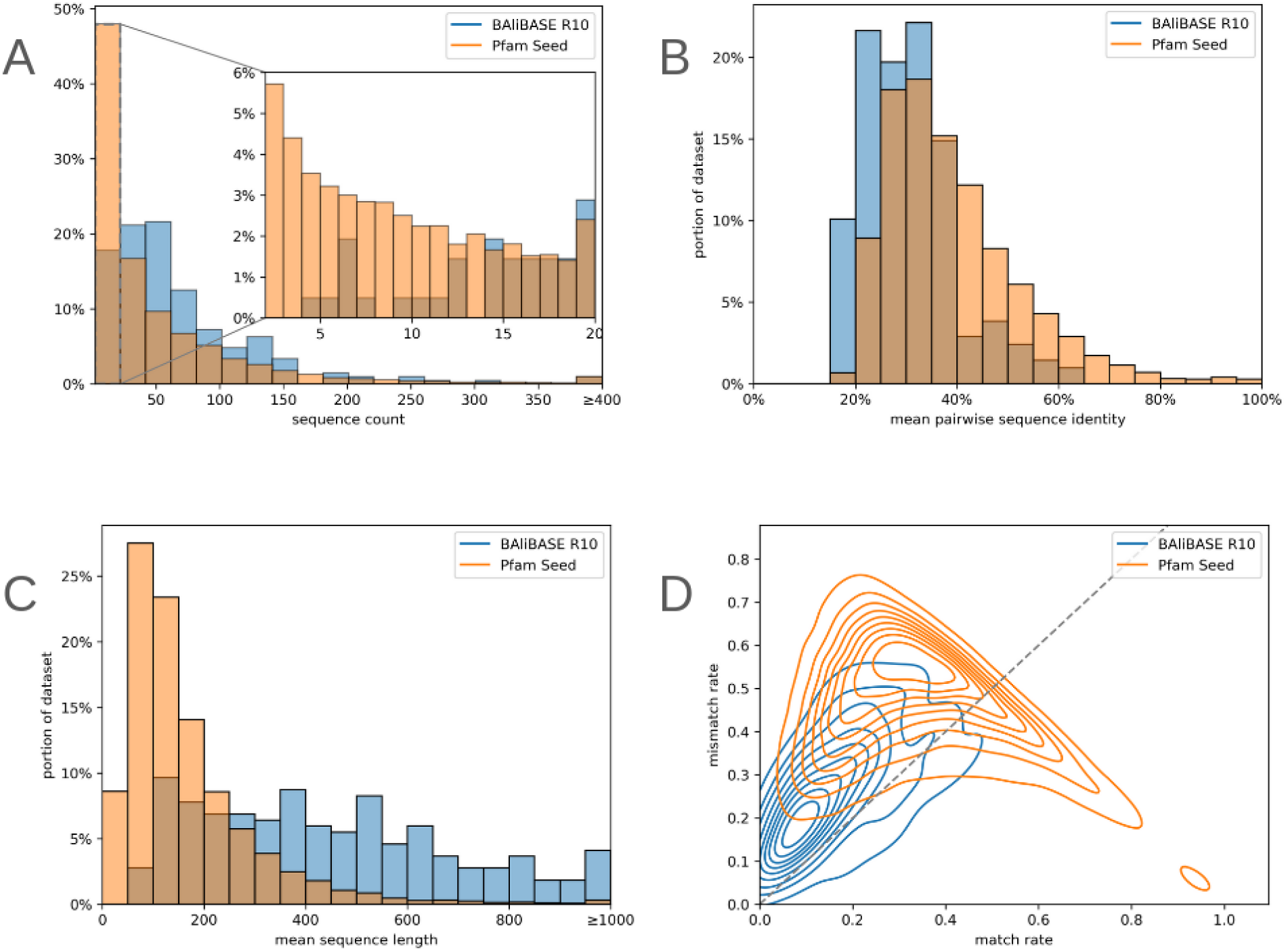
BAliBase vs. Pfam Seed benchmarks in comparison. A, B Distribution of the number of sequences per MSA (A overview for 2 to 150 sequences, B zoomed in for 2 to 20 sequences). C Sequence identity per MSA. D Mean sequence length per MSA. E Match vs. mismatch rate. Match (x-axis) plus mismatch (y-axis) plus gap rate (not shown) equals 100%. Overall (A-E), both BAliBase and Pfam Seed have comparable characteristics as MSA benchmarks, with Pfam Seed MSAs having slightly fewer and shorter sequences of greater sequence identity. Both generally have a higher mismatch than match rate.

One important factor is the number of sequences per MSA (see Figure 3A). BAliBase alignments typically contain a relatively consistent number of sequences, with a mean of 68. In contrast, Pfam displays a distribution closer to a power law, with a mean of 50 but a much lower median of 23. Notably, approximately 6% of Pfam alignments consist of only two sequences, effectively reducing them to pairwise alignments. These cases are considerably easier to align than true MSAs.

Sequence identity within MSAs (Figure 3B) is another relevant factor. We calculated the average pairwise sequence identity across all alignments and observed modest differences: BAliBase alignments averaged 31%, whereas Pfam alignments were slightly higher at 39%. Both are close to the so-called twilight zone (Rost 1999), where sequence comparison becomes difficult. The difference of 8% may be attributable to Pfam’s focus on conserved domains, in contrast to BAliBase’s inclusion of full-length proteins, which often contain more variable regions. As higher sequence identity generally corresponds to easier alignment tasks, this may give a slight advantage to alignments drawn from the Pfam dataset.

Another notable distinction between the two benchmarks lies in sequence length (Figure 3C). Pfam covers domains and BAliBase entire proteins, resulting in longer sequences on average. In addition to sequence length and pairwise sequence identity, we also examined the overall match and mismatch rates across the two benchmark datasets. The match rate is defined as the number of matches divided by the sum of the number of matches, mismatches, and residue-gap pairs. The bivariate density plot of the mismatch and match rates (Figure 3D) indicates that there are relatively more mismatches than matches in both MSA benchmarks. BAliBase is closer to the origin than Pfam Seed because it has more gap-residue pairs. Pfam Seed forms a counter diagonal, which corresponds to ungapped MSAs. Overall, BAliBase alignments exhibit very similar ratios of match to mismatch rates among all residue-residue pairs, suggesting that the underlying alignment difficulty is broadly comparable to Pfam Seed in terms of residue correspondences.

However, differences in absolute match and mismatch rates arise primarily due to varying gap rates. BAliBase alignments tend to include more gaps, resulting in a lower total number of residue–residue pairs. In contrast, Pfam Seed alignments often contain fewer gaps, leading to a higher proportion of residue-residue pairs. This distinction is particularly evident in the density plot of Figure 3E of match versus mismatch rates, where Pfam alignments form a sharp boundary around the line where the sum of match and mismatch rate is one, reflecting the near absence of gaps in many of these alignments.

Besides BAliBase and Pfam Seed as MSA construction tasks, we evaluated the matrices’ performance in three remote homology tasks. The structural domain classification databases CATH and SCOP (Orengo et al. 2003, Murin et al. 2020) classify domains hierarchically into more closely related families and more distantly related superfamilies. The remote homology search tasks consist of determining whether a pair of sequences belongs to the same superfamily or not. The difficulty of this task can be varied by the amount of redundancy per superfamily. Filtering each superfamily at 20% sequence identity (CATH S20 dataset) is more challenging than at 40% (SCOPe40 dataset), and 40% is more challenging than not filtering at all (CATH dataset). For details, see methods.

### Benchmark Results

The sequences taken from the reference MSAs from BAliBase and Pfam Seed are realigned using the method under evaluation. Then the resulting alignments are compared against the original benchmarks. To quantify alignment accuracy, we used two widely accepted metrics (Edgar 2009, Keul et al. 2017, Long et al. 2016):

Q-Score (Sum-of-Pairs Score):

The Q-score evaluates how many residue pairs are correctly aligned across all sequences. It does not require entire columns to be preserved, making it a more permissive and commonly used metric in MSA evaluations.

TC-Score (Total Column Score): This metric represents the percentage of correctly aligned columns relative to the reference MSA. It is a stringent measure, requiring complete agreement across all sequences in a column.

To assess the quality of the three homology search tasks, we report for each the area under the ROC curve.

### AFSM vs. BLOSUM62

Assuming that structural alignments are more accurate than sequence alignments and given that roughly half of the entries in AFSM are different from BLOSUM62, one can expect modest improvements for the evaluation metrics (TC-score and Q-score). This is, however, not the case (see Table 1). AFSM and BLOSUM62 perform nearly identically. This outcome is consistent across the BAliBase (Q-Score 71%/70%) and Pfam Seed (82%/81%) MSA construction benchmarks and the three protein homology search tasks. As expected based on prior comparisons between the benchmarks, alignments on the Pfam Seed dataset yielded slightly better scores overall. This likely reflects the higher sequence identity and lower gap content of domain-focused alignments compared to the more variable and gap-rich alignments in BAliBase. Similarly, the results for AFSM (87%, 79%, 64%) and BLOSUM62 (87%, 80%, 64%) consistently reflect the difficulty of homology search tasks. Interestingly, gap penalties, which were optimised per matrix for the MSA construction task, were remarkably similar (see Table 1).

### Choice of substitution matrices

The above results are unexpected. Despite more and other data, AFSM and BLOSUM62 perform nearly identically across five different tasks. To put this finding into perspective, we added 16 other matrices to the analysis, including classical sequence-based series PAM and BLOSUM (Henikoff et al. 1992), more recent improvements such as VTML (Müller et al. 2000), RBLOSUM, and CORBLOSUM (Hamacher et al. 2016), as well as structure-derived matrices like SDM and HSDM (Prlić et al. 2000), ProtSub (Jia et al. 2021), and PFASUM (Keul et al. 2017). Besides these established matrices, we added two controls, which only distinguish matches and mismatches. To make these controls comparable to the above matrices, we assigned matches a score of 5, which is in line with the other matrices whose diagonals average to values of 4 to 6. For mismatches, we considered two scores of 0 and -1, respectively. Essentially, these two controls reflect the absence of a substitution matrix, and since they are similar to a unit matrix, we named them baseline unit matrix 5/0 and 5/-1.

### Growth of data

Table 1 lists the number of residue pairs used in the calculation of each substitution matrix. Over the decades spanning the development of these matrices, there has been an increase of roughly two orders of magnitude in the amount of data used. For example, the original PAM matrices, introduced in the late 1970s, were based on an estimated 470,000 residue pairs. In this work, we derived the AFSM from over 68 million residue pairs, reflecting both the growth in available data and advances in computational methods.

### Similarity of substitution matrices

To provide an overall impression of the similarity between matrices, we represented each substitution matrix as a high-dimensional vector and applied dimensionality reduction, as shown in Figure 1B. The resulting projection reveals that most matrices—including the BLOSUM series, AFSM, Gonnet, VTML, and PFASUM matrices—cluster closely together near the center, reflecting their high degree of similarity. In contrast, the baseline matrices form a distinct cluster separate from this main group, as do the PAM matrices. The structure-derived SDM and HSDM matrices also lie outside the central cluster. They are separated from each other as well.

### Nearly all substitution matrices are similar to BLOSUM62

Despite differences in approach—whether the matrices were derived from sequence alignments or structural data, as indicated in Table 1—and substantial variation in the amount of data used for their construction, the matrices show remarkably little deviation from BLOSUM62. Table 1 includes a column reporting the correlation of each matrix with BLOSUM62. Most matrices (excluding the baseline unit matrices) exhibit a correlation above 84%, with the majority clustering even higher, at 95% or above. This highlights the overall similarity of substitution patterns captured by these matrices, regardless of their underlying methodology or data scale.

### Parameter Optimization

To ensure a fair comparison of the matrices, we individually optimized the gap opening and extension penalties for each matrix and benchmark. Table 1 summarizes these settings, showing that gap opening penalties ranged from 4 to 9, while extension penalties were consistently low, with most values at 0.5 across all configurations. Figure 4 gives an impression of the change in Q-score of the AFSM matrix as gap parameters vary. For gap opening penalties ranging from 0 to 9 and gap extension ranging from 0 to 4, one sees a relatively consistent Q-score for most parameter combinations greater than 2 for opening and around 0.5 and 1 for extension. The optima are close to each other for both Pfam Seed and BAliBase, but they are not identical.

**Figure 4.**
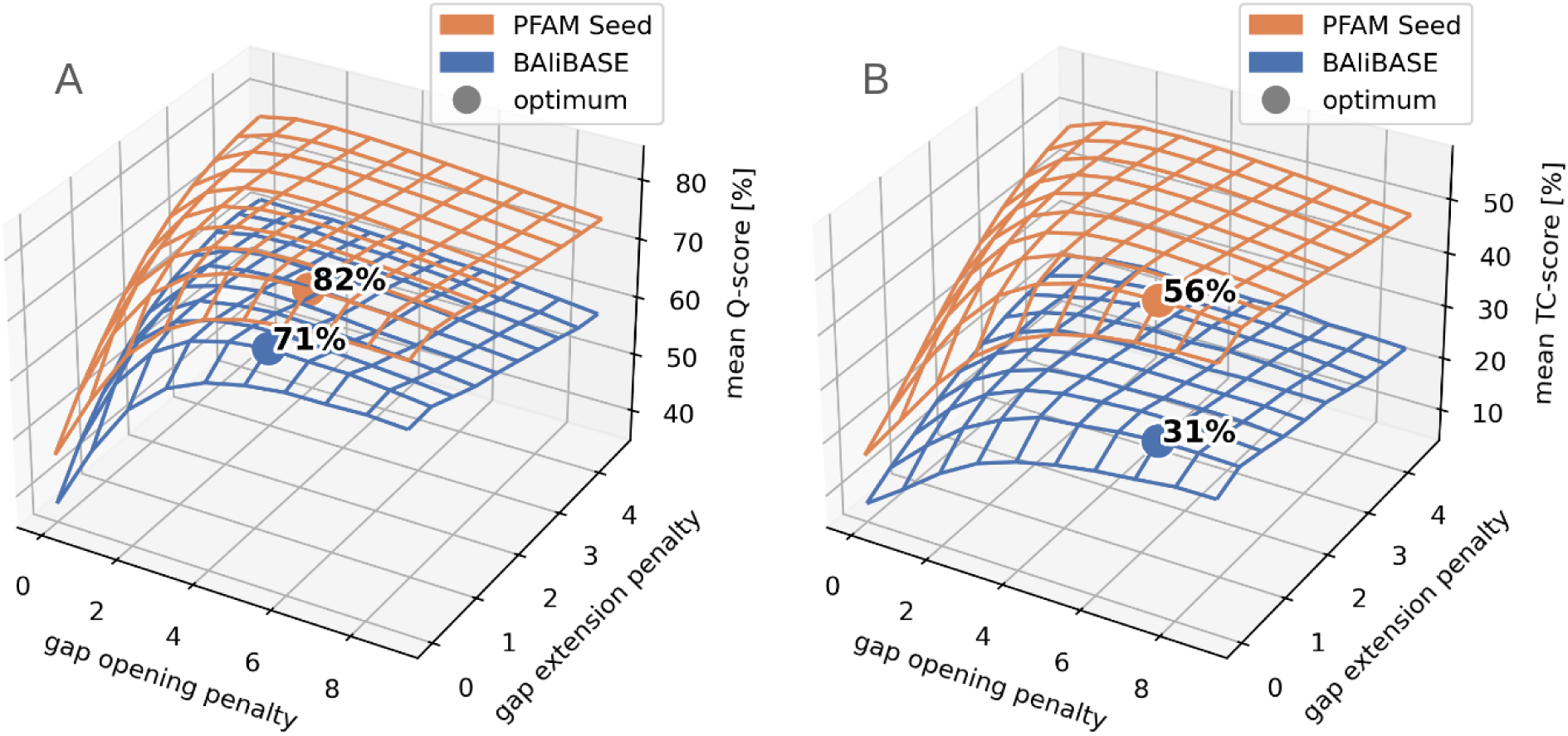
Optimal gap penalties and corresponding alignment scores for the AFSM matrix on BAliBASE and Pfam Seed benchmarks. (A) Q-scores and (B) TC-scores across a range of gap opening (GOP) and gap extension (GEP) penalties. For BAliBASE, the highest Q-score (71%) was achieved with GOP 5 and GEP 0.5, while the highest TC-score (31%) was obtained with GOP 7 and GEP 0.5. For Pfam Seed, the optimal Q-score was 82% with GOP 6 and GEP 0.5, and the highest TC-score was 56% with GOP 7 and GEP 0.5. Both benchmarks show similar trends and relatively stable scores across a broad range of parameter settings.

### Nearly all substitution matrices perform similarly to BLOSUM62

Despite differences in origin, age, and the amount of data used, the results reveal a striking consistency: most matrices designed for low sequence identity scenarios perform nearly identically (see Table 1). Q-scores were consistently higher on Pfam Seed alignments (81–83%) compared to BAliBase (69–73%), and the same trend was observed for TC-scores (54–57% for Seed; 30–33% for BAliBase). Notably, corrected matrices derived from BLOSUM (RBLOSUM, CorBLOSUM) did not lead to measurable performance gains, consistent with findings from previous studies (Hamacher et al. 2016). Furthermore, no clear relationship was observed between alignment performance and the number of residue pairs, year of matrix publication, or the type of data used (sequence-vs. structure-based). Similarly, there were no strong performance deviations for the homology search tasks. Most of the matrices perform with a maximum 2% difference from BLOSUM62 for the ROC AUC and similarly for the PR AUC curves. All matrices consistently show that homology search is more difficult when redundancy is low. Interestingly, the baseline unit matrix 5/0 is the single matrix performing significantly worse than BLOSUM62 in the homology search tasks, despite performing equally well in MSA construction.

### BLOSUM80 and PAM120 perform worse than BLOSUM62

BLOSUM80 and PAM120 are designed to compare highly similar sequences rather than remote homologues. Consequently, these two matrices perform significantly worse than BLOSUM62. There is one interesting exception: For the most challenging CATH S20 homology search task, BLOSUM80 performs 5% better than BLOSUM62.

### Baseline unit matrices perform as good as BLOSUM62

A striking result of our study was the performance of the baseline unit matrices. These simple matrices consist of a single conservation score on the main diagonal and a single mismatch score for all off-diagonal entries. To match the average conservation signal of BLOSUM62, we set the conservation score to 5, which resulted in similar optimal gap penalties to those found for BLOSUM62. We tested two versions of the matrix: one where mismatches were ignored (mismatch score 0), representing a matrix suited for high evolutionary distances, and one where mismatches were mildly penalized (mismatch score −1), representing a matrix suited for low evolutionary distances like BLOSUM80 or PAM120.

Contrary to our expectations, both baseline matrices performed surprisingly well. Each performed on a par with their corresponding established matrices. Remarkably, the baseline matrix 5/0 that ignored mismatches even slightly outperformed BLOSUM62 on BAliBase, achieving Q-scores that were higher by about one percentage point. Similarly, baseline matrix 5/-1 performs close to BLOSUM8. Regarding the protein homology search tasks, the no matrix options generally perform slightly worse than BLOSUM62. However, the baseline unit matrix 5/-1 even outperforms BLOSUM62 on the most challenging CATH S20 task. This indicates that gap penalties play an equally important role as match/mismatch scores.

In summary, these findings underscore that the specific choice of substitution matrix has only a limited impact on MSA construction and homology search, provided the matrix is broadly appropriate for the evolutionary context.

### Performance on MSA construction is dependent on MSA size

The good performance of the baseline unit matrices is surprising. One explanation may be that substitution matrices are useful when there are few sequences at a moderate distance. But as there are more and more sequences in an MSA, they impose more and more constraints arising from matches, which make the alignment task easier and not more difficult. To test whether this could be true, we plotted the improvement in Q- and TC-scores for BLOSUM62 and AFSM over the Baseline Unit Matrix 5/0 against the number of sequences in the MSA. Figure 5 shows that the Q- and TC-scores for BLOSUM62 are slightly worse than using no matrix whenever there are more than around 20 sequences in the alignment. For very small numbers of sequences, BLOSUM62 improves an MSA, but for larger ones, it slightly deteriorates an MSA. This implies that BLOSUM62 was beneficial when there was little sequence data available, but its usefulness is reduced as sequence data grows.

**Figure 5.**
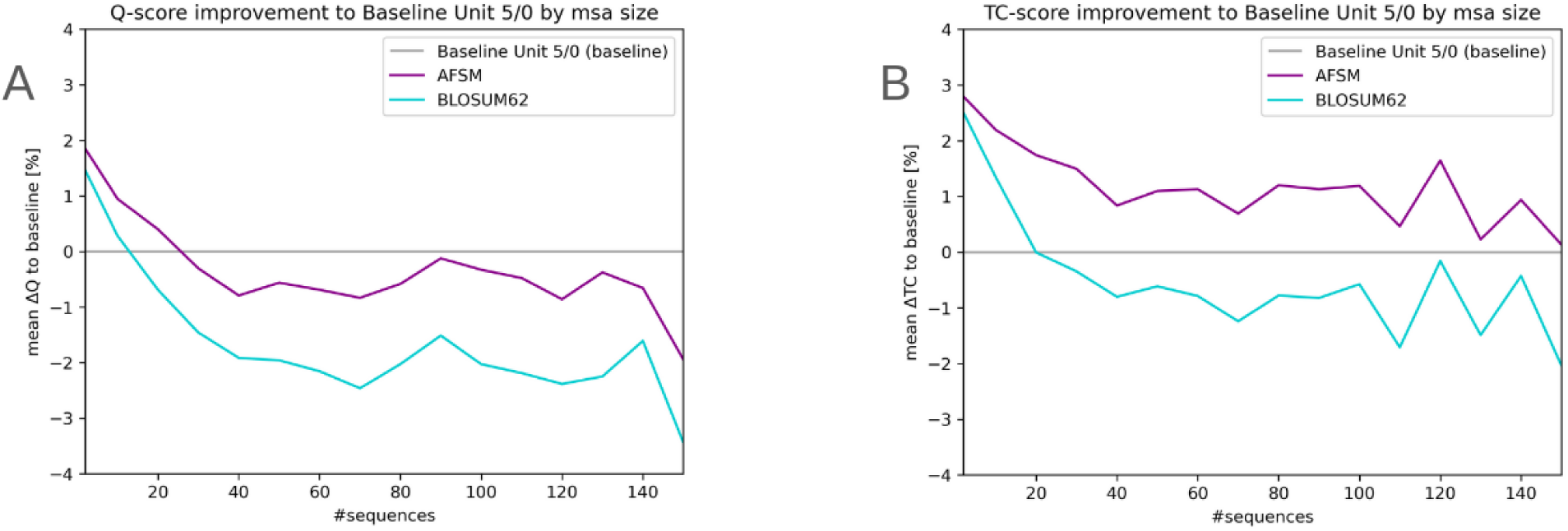
Q-score (A,C) and TC-score (B,D) of AFSM and BLSOUM62 for Pfam Seed compared to a baseline of using “no matrix” (Baseline Unit Matrix 5/0), in dependence of the number of sequences in an MSA. A, B overview for up to 150 sequences, Generally, the more sequences there are, the lower the difference from the baseline. AFSM performs slightly better than BLOSUM62. Most importantly, for small amounts of sequences, AFSM and BLOSUM62 are better than no matrix; for larger amounts of sequences, they are worse.

### Embeddings surpass substitution matrices when sequence data is sparse

Taken together, the results of this study are intriguing. Independent of the amount of data and the type of data used, matrices perform comparably. And even the baseline unit matrices, which essentially amount to using no matrix at all, perform on a par. Does this mean that the vast amounts of sequence and structure data available today make no difference? To put this in context, we performed the homology search task on embeddings. Embeddings (Rost et al. 2022) are numerical vectors representing sequences. They result from training a neural network on billions of sequences, thus capturing the essence of sequence space in a compact form.

Embeddings can be treated as numerical vectors, which can be compared by Euclidean distance. For each of the sequences in the three homology search datasets, we obtained its embedding and selected the domain with the closest embedding (see methods), i.e. we implemented a nearest neighbor search. We then evaluated whether the query domain and its nearest neighbor are in the same superfamily. From this we computed an AUC. Similarly, we obtained for the query the domain with the highest similarity score using AFSM and BLOSUM62. And from this we computed the AUC in an analogous manner.

We found that embeddings achieve an area under the ROC for the three homology search tasks of 99% (CATH), 94% (SCOPe40), and 90% (CATH S20). The differences nicely reflect the difficulty of the data sets resulting from the increase in sparsity. In comparison, BLOSUM62 and AFSM achieve both 99%, 89%, and 78%, respectively and the baseline unit matrices 5/-1 and 5/0 99%, 85%, 74% and 99%, 57%, and 62%. This is a far-reaching result. It means that when sequence data is dense, classical sequence search with or without substitution matrices performs as good as embeddings do. However, when sequence data is sparse, embeddings make a difference. Having implicitly absorbed billions of sequences, embeddings perform 12% better than the two substitution matrices. Interestingly, for the sparse data set, one of the baseline matrices performs as good as BLOSUM62 and AFSM. Overall, this means that embeddings are very valuable where large evolutionary distances have to be covered, i.e. when comparing e.g. protein sequences from a newly sequenced genome to model organism sequences. But, it also means as sequence data is growing and as the overall sequence density saturates, classical sequence alignment with or without substitution matrices performs comparatively.

## Discussion and Conclusion

BLOSUM62 was introduced over 30 years ago. Although it was corrected, recalculated on larger sequence data, complemented by different statistical approaches and by use of structural instead of sequence data, it nonetheless remains a default substitution matrix in protein BLAST searches. Why is BLOSUM62 still in use? Our results may contribute to an explanation. None of the existing approaches presented in this article, including AFSM, substantially improve MSAs or homology search in comparison to BLOSUM62. Logically, alternatives are not widely adopted. But in this article, we showed that BLOSUM62 appears not to perform better than using no matrix on the used benchmarks. In fact, only when very few sequences are used, it performs slightly better than using no matrix. These results suggest that BLOSUM62 was important in the early days of sequence databases, when data was sparse. But in today’s world of billions of protein sequences, our results suggest that BLOSUM62 is of limited use. This is further supported by our results on the use of embeddings in the three homology search tasks. Embeddings are statistical representations of sequences, incorporating the total population of all sequences. Thus, embeddings can be assumed to implicitly include knowledge of observed substitution patterns, which is made explicit in substitution matrices. The performance gap between embeddings and sequence comparison with a substitution matrix is large (12%) so that homology search should consider the use of embeddings. It is not obvious how embeddings can be used to construct MSAs, but (36734516) introduces a deep neural network, which proves that language models can be adopted for this task. Another alternative may be novel representations of sequences. Edgar introduces a mega-alphabet, on which classical sequence algorithms perform as good as structure-based approaches (Edgar 2024). An intermediate approach using embedding-based protein sequence alignments is proposed by Schwede et al. (Schwede et al. 2024). Do all of these considerations mean that there is no need for BLOSUM62 and other matrices like AFSM? While substitution matrices including AFSM appear not to improve MSA construction and protein homology search, they also do not make results worse, and they turn the intuition of chemists and physicists into numbers. Indeed, this may be one of the main reasons for BLSOUM62’s popularity. Manually assigning the physicochemical properties of amino acids numerical values is difficult (though there were attempts in the past (Granham 1974)). Moreover, in pairwise alignments or areas where sequence data is sparse, substitution matrices will still be needed to bridge the gaps between evolutionarily distant sequences.

## Methods

Pfam Seed release 35 was accessed from ftp.ebi.ac.uk in March 2024. After filtering out large MSAs (>100KB), 18,061 alignments with a total of 909,284 sequences were left. BAliBase release 10 (R10) was accessed from www.lbgi.fr in February 2024. CATH version 4.3.0 and SCOPe version 2.07 were used. CATH S20 and SCOPe40 datasets were not filtered. Due to size, CATH was reduced by removing very short (<50AA) and very long (>250AA) sequences, resulting in 23,911 domains. Interpro release 95 was accessed from ftp.ebi.ac.uk. The code to calculate Q- and TC-scores was obtained from drive5.com/qscore.

AFSM. To construct AFSM, 297 domain families with up to 100 domains each were randomly selected from Interpro. Not all families have 100 domains. To balance the difference in domain numbers per family, we introduce a weighting factor w = 2 / (k(k−1)), where k is the number of domains for a family, i.e., w relates to the number of domain pairs within a family. Structures for all domains across all families were obtained from the AlphaFold database (accessed in March 2024), where available. This resulted in 20.842 structures. All pairs of domains with less than 62% sequence identity (obtained with Needleman-Wunsch algorithm with match 1, gap -1, mismatch -1) were structurally aligned using PyMOL’s ce-align. Alignments with a poor RSMD value of >7Å were discarded.

AFSM scores were computed following the BLOSUM methodology, with slight deviations in how substitution frequencies were acquired:

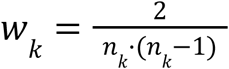

The weighting factor for family *k*, containing *n_k_* member proteins.

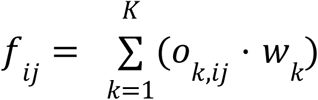

Weighted substitution frequency of amino acids *i* and *j*, where *o_k,ij_* is the number of observed substitutions between *i* and *j* in family *k*, and *K* is the total number of families.

From this point, we followed the BLOSUM methodology: substitution odds were calculated by dividing observed substitution frequencies by expected substitution frequencies, taking the base-2 logarithm of the result, multiplying by two to express scores in half-bit units, and rounding to the nearest integer.

In Table 1, Corr reports Pearson correlation (corrcoef in NumPy) and Evo Dist is mean diagonal (conservation) by mean off-diagonal (mutation).

Sequence identity in Figure 3B was computed with parasail NW with gap open=10, gap extend=1, matrix=blosum62.

Match and mismatch rates in Figure 3E were calculated per MSA in the benchmark as the proportion of identical (match) and non-identical (mismatch) residue–residue pairs, relative to the total number of residue–residue and residue–gap pairs.

MSAs were computed for the benchmarks and substitution matrices using ClustalW2 version 2.1 downloaded from clustal.org/download in March 2024.

AUC. For all matrices and all benchmarks, the Area Under the Curve (AUC) of the Receiver Operating Characteristic (ROC) curve was calculated using scikit-learn’s roc_auc_score function (Version 1.6.1).

For homology search, sequence alignment scores were obtained with parasail NW with optimal gap open and extension penalties as shown in Table 1. As parasail only accepts integers and optimal gap penalties were sometimes floating-point values, substitution matrices and gap penalties were scaled up accordingly. For AUC of homology search, similarity of all pairs of protein sequences of the Cath, Cath S20, SCOPE40 datasets were recorded separately for pairs in the same and in different superfamilies. From this AUC curves were computed.

Embeddings were obtained using ProtTransT5XLU50 v0.2.2. (<u>github.com/sacdallago/bio_embeddings</u> Rost et al. 2022). To obtain the nearest neighbor of each embedding, the KNN-library FAISS (version 1.8.0) was used with Euclidean distance (L2) implemented by IndexFlatL2. After index construction, KNN searches were parallelized across queries using the joblib library (version 1.4.2) to enhance performance on a compute cluster.

AUC was computed by binarizing the labels and applying the one-vs-rest (OvR) approach from the sklearn.metrics module (version 1.3.2) with average=’micro’ and multi_class=’ovr’.

Convergence Analysis

To determine how data size influences the AFSM values, we varied the number *n* of families from 1 to 297 used to construct an AFSM. For each *n*, we repeated the analysis a 1000 times. In Figure 2B we report the absolute position-wide difference between the full and partial AFSM.

### Residue Pair Estimation

To assess the size of the data used for construction of the substitution matrices (Table 1), we estimated the number of residues pairs as follows:

PAM1 is based on 1,572 mutations, which leads to 471,000 (=1,572*300) residue pairs assuming an average sequence length of 300 (book.bionumbers.org/how-big-is-the-average-protein). For PFASUM, we counted the total number of residue pairs in the full set of Pfam Seed alignments at around 584 million. Since PFASUM applies the same clustering strategy as BLOSUM, we assume that it retains a similar fraction of residue pairs, allowing us to approximate the number of pairs used in each matrix variant based on known BLOSUM yields.

### Benchmarking Methods

Gap opening and extension penalties were determined by exhaustive search (0-10 and 0-5, respectively). Benchmark MSAs were stripped of gap characters and realigned using ClustalW2 with the following options: -type=protein, -output=fasta, -negative, -matrix=[…], -gapopen=[…], -gapext=[…]. The resulting alignments were compared to the original benchmark using qscore. The full benchmarking pipeline was implemented using Nextflow.

## Acknowledgement

Writing was assisted using ChatGPT (GPI-4o) and Grammarly (v 9.63). Neither tools were used for text generation from scratch but solely for correction in spelling and grammar. All suggested changes from AI tools have been carefully checked for correctness of content. Funding by BMBF projects Ebira and scads.ai is acknowledged. Thanks for technical support to Alexandre Mestiashvili. Special thanks to Ali Al-Fatlawi for cleaned SCOP and CATH datasets.

